# Coherent pathway enrichment estimation by modeling inter-pathway dependencies using regularized regression

**DOI:** 10.1101/2022.07.06.498967

**Authors:** Kim Philipp Jablonski, Niko Beerenwinkel

## Abstract

Gene set enrichment methods are a common tool to improve the interpretability of gene lists as obtained, for example, from differential gene expression analyses. They are based on computing whether dysregulated genes are located in certain biological pathways more often than expected by chance. Gene set enrichment tools rely on pre-existing pathway databases such as KEGG, Reactome, or the Gene Ontology. These databases are increasing in size and in the number of redundancies between pathways, which complicates the statistical enrichment computation. Here, we address this problem and develop a novel gene set enrichment method, called *pareg*, which is based on a regularized generalized linear model and directly incorporates dependencies between gene sets related to certain biological functions, for example, due to shared genes, in the enrichment computation. We show that *pareg* is more robust to noise than competing methods. Additionally, we demonstrate the ability of our method to recover known pathways as well as to suggest novel treatment targets in an exploratory analysis using breast cancer samples from TCGA. *pareg* is freely available as an R package on Bioconductor (https://bioconductor.org/packages/release/bioc/html/pareg.html) as well as on https://github.com/cbg-ethz/pareg. The GitHub repository also contains the Snakemake workflows needed to reproduce all results presented here.

## 1 Introduction

The behavior of cells is governed by a complex interplay of molecules. Their functional dynamics are organized according to biological pathways (Chuang *et al*., 2010). Perturbations of pathways have been linked to certain diseases, such as, for example, cancer (Hanahan and Weinberg, 2000, 2011). Biological pathways can be obtained from pathway databases such as the Kyoto Encyclopedia of Genes and Genomes (KEGG), the Gene Ontology (GO), or Reactome (Ogata *et al*., 1999; Consortium, 2004; Joshi-Tope *et al*., 2005). It is important to note that pathways typically impose a structure of interactions in the form of a network on its contained molecules. While the nodes of this network typically correspond to genes, the edges correspond to interactions, such as signal transductions (Steffen *et al*., 2002). Another way of grouping genes in a meaningful way is to forgo the structure requirement and simply consider, for example, functionally related genes to be part of the same gene set.

Experiments investigating, for instance, differentially expressed genes between several conditions (e.g., wild-type versus mutant cell cultures) often produce a long list of genes of interest which is difficult to interpret (Simillion *et al*., 2017; Maleki *et al*., 2020). A common method for aggregating these lists of potentially interesting genes is to assess whether the genes preferentially appear in biologically relevant pathways. This reduces the amount of information which needs to be interpreted from individual genes to groups of genes, i.e., pathways, following a similar function.

There are several approaches to computing whether certain genes preferentially appear in certain gene sets. They can be roughly divided into the three groups: (a) singular enrichment analysis, (b) gene set enrichment analysis, and (c) modular enrichment analysis (Huang *et al*., 2009). In a singular enrichment analysis, a list of genes resulting from a differential expression analysis is first partitioned into differentially expressed and not differentially expressed genes based on a threshold typically applied to effect size or p-value. These two groups of genes are then used to compute a pathway enrichment score individually. The gene set enrichment analysis lifts the requirement of a pre-selection of genes and considers all input genes without partitioning them into groups based on a threshold. Finally, the modular enrichment analysis computes the enrichment of each gene set not in isolation but rather by incorporating term-term relations into the statistical model. A term is a set of genes which are all involved in the same biological process and are thus functionally related. These term-term relations represent dependencies between gene sets, which can arise, for example, due to shared genes. This approach has the advantage of not requiring arbitrary thresholds to prepare the input genes and is able to incorporate additional biological knowledge into the enrichment computation by imposing a structure on the gene set database. This additional biological knowledge can help maintain high statistical power in large, redundant gene set databases or structure the final visual presentation of enrichment scores (Huang *et al*., 2009).

One of the most basic approaches to compute singular enrichments is to use Fisher’s exact test which is based on the hypergeometric distribution and requires a stratification of the input gene set (Fisher, 1922). There have been many extensions to this initial approach, including threshold-free methods such as the popular tool GSEA (Subramanian *et al*., 2005) which does not require an a priori stratification of the input and LR-Path which formulates the enrichment computation as a regression (Sartor *et al*., 2009).

Various methods have been proposed which follow the modular enrichment approach. topGO (Alexa *et al*., 2006) is tailored to the tree structure of the gene sets provided by the Gene Ontology resource and removes local dependencies between GO terms which leads to better performance. By relying on the topology of a tree, it is not applicable to many other gene set sources. Another approach is to reduce the number of pathways which are included in the enrichment computation by removing redundant terms based on the notion of semantic similarity (Yu *et al*., 2010; Wu *et al*., 2021). RedundancyMiner (Zeeberg *et al*., 2011) transforms the GO database prior to the enrichment computation by de-replicating redundant GO categories and thus tries to reduce the amount of noise introduced by overlapping pathways appearing in the enrichment analysis. These approaches rely on the directed acyclic graph structure of GO terms and cannot be generalized to other pathway databases. GENECODIS (Carmona-Saez *et al*., 2007) incorporates relations between pathways into the enrichment computation by testing for the enrichment of co-occurring pathways. It can in principle be applied to any pathway database but is only available as a web-based tool and can thus not be easily used in automated workflows. The same limitation applies to ProfCom (Antonov *et al*., 2008) which computes the enrichment of unions, intersections, and differences of pathways. In addition, it uses a greedy heuristic which does not guarantee to find an optimal solution for each case. MGSA (Bauer *et al*., 2010) embeds all pathways in a Bayesian network and identifies enriched pathways using probabilistic inference. It does however not allow to explicitly model pathway relations. Finally, tools such as EnrichmentMap (Merico *et al*., 2010), ClueGO (Bindea *et al*., 2009), REVIGO (Supek *et al*., 2011) and GOrilla (Eden *et al*., 2009) compute a singular enrichment score per pathway and subsequently visualize the result as a network of gene set clusters based on gene overlaps. This approach can be applied to any gene set database but loses statistical power by executing the enrichment analysis and term-term relation inclusion in separate steps.

While many methods exist which try to overcome the issue of large redundant pathway databases, none of them, to the best of our knowledge, has accomplished this goal in a simultaneously database-agnostic, flexible and robust way. By not relying on the hierarchical structure of the Gene Ontology it is possible to create a method which is less restricted and can be used with other pathway databases that are more specialized to the experiment at hand. As there are various approaches to comparing pathways with each other, it is desirable for the enrichment algorithm to not be hard-coded to use a single specific pathway similarity measure but allow different ones based on the needs of the respective research question. The noise inherent to biological experiments leads to measurements of differential gene expression which can deviate from the underlying true differences. Robustness to the level of noise of the input data is thus a crucial property of pathway enrichment methods.

Here, we introduce a novel method called *pareg* for computing pathway enrichments which is based on regularized regression. It follows the ideas of GSEA as it requires no stratification of the input gene list, of MGSA as it incorporates term-term relations in a database-agnostic way, and of LRPath as it makes use of the flexibility of the regression approach. By regressing the differential expression p-values of genes on their membership to multiple gene sets while using LASSO and gene set similarity-based regularization terms, we require no prior thresholding and incorporate term-term relations into the enrichment computation. We show in a synthetic benchmark that this model is more robust to noise than competing methods, and demonstrate in an application to real data from The Cancer Genome Atlas (TCGA) (Tomczak *et al*., 2015) that it is able to recover known pathway associations as well as suggest novel ones.

## 2 Methods

### Overview

The input to *pareg* consists of (a) a list of genes, where each gene is associated to a single p-value obtained from a differential expression experiment and (b) a gene set database where a gene can be part of multiple gene sets simultaneously. *pareg* ‘s approach is general enough to support any kind of experimental value associated to the input genes. Pathway enrichments are then computed by regressing the differential expression p-value vector of input genes on a binary matrix indicating gene membership to each gene set in the input database. The estimated coefficient vector captures the degree of association which gene sets have with p-values of differentially expressed genes; they can thus be regarded as an enrichment score. To induce sparsity in the coefficient vector and thus in the selected set of enriched pathways, we use the least absolute shrinkage and selection operator (LASSO) regularization term (Tibshirani, 1996). Term-term relations are included in the model using a network fusion penalty (Cheng *et al*., 2014; Dirmeier *et al*., 2018).

### Regression approach

We use a regularized multiple linear regression model to estimate gene set enrichment scores. Suppose we want to compute the enrichment of *K* pathways using *N* genes. Each gene *g*_*i*_ is associated with a p-value *p*_*i*_ from a differential expression analysis for *i* = 1, …, *N*. We then define the response vector **Y** to be

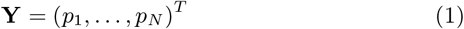

The binary regressor matrix **X** captures the membership information of each gene *g*_*i*_, *i* = 1, …, *N*, in pathway *t*_*j*_, *j* = 1, …, *K*,

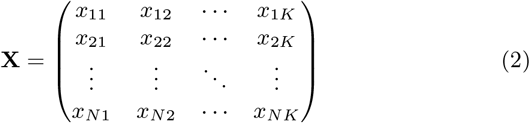

with

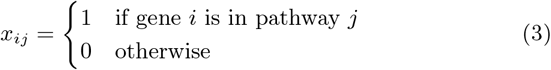

In the resulting linear model **Y** = **X***β*, the vector of coefficients *β* = (*β*_1_, …, *β*_*K*_)^*T*^ is estimated using stochastic gradient descent to minimize the objective function

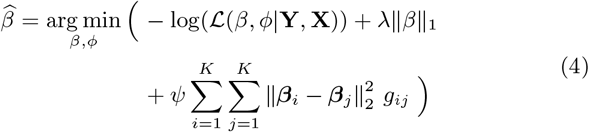

where *ℒ* (*β, ϕ*|**Y, X**) is the likelihood and **G** = (*g*_*ij*_)_*ij*_ *∈* (0, 1)^*K×K*^ a pathway similarity matrix, where *g*_*ij*_ describes the similarity between pathway *i* and *j*.

To model the p-values in the response vector, the likelihood is defined using the beta distribution (Ferrari and Cribari-Neto, 2004)

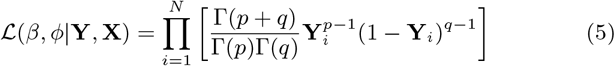

where *p* = *µϕ* and *q* = (1 *−µ*)*ϕ* with mean 0 *< µ <* 1, precision parameter *ϕ >* 0 and Gamma function Γ(·). The mean is then modeled as *g*(*µ*) = **X***β* where *g*(·) is a link function (Cribari-Neto and Zeileis, 2010).

The optimal values for the regularization parameters *λ* (LASSO) and *ψ*(network fusion) are determined using cross-validation (Dirmeier *et al*., 2018), which balances the effects of the LASSO and network fusion terms. The former term induces a sparse coefficient vector, i.e., it reduces the number of enriched pathways needed to explain the observed data. The latter term promotes assigning a similar enrichment score to (functionally) similar pathways.

### Pathway similarity measures

The goal of adding pathway similarities to the model is to group pathways in the enrichment computation. By doing so, redundant sets of functionally related pathways jointly drive the enrichment signal and reduce the influence of noisy measurements. Due to the flexibility of our model, this can be any similarity measure which can be stored as a real matrix.

As pathways are typically defined as lists of genes, the Jaccard similarity and overlap coefficients are common choices (Merico *et al*., 2010). They group pathways which share many genes together and are thus a good measure of functional relation (Bass *et al*., 2013). The overlap coefficient is particularly suited for pathway collections which feature a hierarchical structure.

In addition, when using the popular Gene Ontology (Consortium, 2004) as a pathway database, semantic similarity measures exist. These measures incorporate the topological structure of the Gene Ontology and are better at inferring functional relations between pathways (Guo *et al*., 2006; Ehsani and Drabløs, 2016; Zhao and Wang, 2018).

### Presentation of enrichment results

The estimated coefficient vector *β* can be ordered descendingly by absolute value such that the most dysregulated and thus interesting pathways appear at the top of the list. A regression coefficient *β*_*j*_ of large absolute value corresponds to a strong dysregulation of pathway *j*.

In addition, we implement a network-based visualization of the enrichment result. Each node in this network corresponds to a pathway, and edges correspond to high pathway similarities. The nodes are colored by the respective enrichment score of each pathway. This allows for the quick identification of functional modules as network clusters.

Finally, the result of *pareg* can be transformed to a format readily understood by the functional enrichment visualization R package enrichplot (Yu, 2022). This enables the usage of many plotting functions, such as dot plots, tree plots and UpSet plots, as well as immediate access to newly implemented ones.

### Generation of synthetic data

The goal of the synthetic benchmark is to create a known set of dysregulated pathways which induces a set of differentially expressed genes, apply several enrichment methods (listed below) to this data set and evaluate how well each method is able to recover the initially dysregulated pathways. Thus, each synthetic data set consists of a list of genes with associated p-values obtained from a simulated differential expression experiment, as well as a respective ground truth set of pathways.

Given an existing term database *D* = *{T*_1_, …, *T*_*K*_ *}* consisting of *K* terms 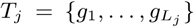, each made up of *L*_*j*_ genes *g*_*i*_, we randomly sample a ground truth set of activated terms *D*_*A*_ *⊂ D*. In order to model the joint activation of functionally related pathways, we apply a similarity sampling approach. Given a similarity matrix *S* with 0 *≤ s*_*ij*_ *≤* 1 and similarity factor 0 *≤ ρ ≤* 1 we first uniformly sample a single term *j*. The next term is then drawn according to the probability vector (1*−ρ*)*U* +*ρS*_*j*_ where *S*_*j*_ is column *j* of *S* and denotes the similarity of term *j* to all other terms, and *U* is a vector of length |*S*_*j*_| with values 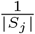. This procedure is continued by setting *j* to the previously sampled term and repeated until the required number of terms has been sampled. For *ρ* close to 1 this results in similar pathways being sampled while *ρ* close to 0 leads to a uniformly random sample.

Next, we model synthetic differential expression p-values for the *N* genes (*g*_1_, …, *g*_*N*_) by sampling from a Beta distribution whose parameters are determined from a linear combination of a noisy gene-term membership matrix and a term activation vector. This mimics the real life setting where the dysregulation of a pathway is jointly driven by the dysregulated genes it contains.

In particular, we create the activation vector *β*_*A*_ = (*b*_1_, …, *b*_*K*_)^*T*^ with

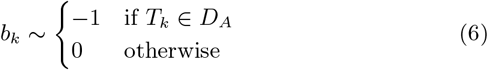

That is, we assign a non-zero coefficient to activated pathways. The gene-term membership matrix **X**_*A*_ is defined analogously to eqs. (2) and (3). To model the effect of noisy measurements, we remove the association between genes and activated terms in **X**_*A*_ by setting a fraction of *η* entries to 0. Next, we compute *µ* = *g*^*−*1^ (**X**_*A*_*β*_*A*_) where *g*^*−*1^ is the logistic function and set *ϕ* = 1 to parametrize the Beta distribution. To create the final synthetic data set *E* = (*D*_*A*_, *{*(*g*_*i*_, *p*_*i*_), …, (*g*_*N*_, *p*_*N*_)*}*), we sample the differential expression p-value *p*_*i*_ for gene *i* from *B*(*µ*_*i*_, *ϕ*).

We run 20 replicates with 20 activated terms each and use all pathways with sizes between 50 and 500 in the biological process subtree of the Gene Ontology.

### Performance evaluation in synthetic benchmark

Due to the strong class imbalance in the experimental setup of pathway enrichments featuring few positives, i.e., dysregulated pathways, compared to the number of negatives, i.e., unaffected pathways, we use precision-recall (PR) curves to evaluate the performance of each pathway enrichment method (Davis and Goadrich, 2006; Saito and Rehmsmeier, 2015).

A term *T*_*j*_ is classified as a true positive (TP) if it is in *D*_*A*_ and is enriched according to a method and respective threshold. It is classified as a false positive (FP) if it is not a member of *D*_*A*_ but is estimated to be enriched. Analogously, a true negative (TN) is a term which is not in *D*_*A*_ and is not enriched, while a false negative (FN) is a term which is in *D*_*A*_ but is not detected by a method. Precision is then defined as *TP/*(*TP* + *FP*) and recall as *TP/*(*TP* + *FN*). By varying the threshold used to create the classifications we can then readily create PR curves. To obtain a numeric summary of a method’s performance, we compute the area under the precision-recall curve (AUC).

### Real data application

We conduct an exploratory analysis using cancer and normal samples from processed TCGA data available in the Gene Expression Omnibus entry GSE62944 (Rahman *et al*., 2015). We retrieve 113 tumor and matched normal samples for TCGA-BRCA (Breast Invasive Carcinoma). We then use limma (Ritchie *et al*., 2015) to run a differential gene expression analysis to compare tumor and normal samples. The obtained p-values and pathways from the biological process subtree of the Gene Ontology are then used as input to *pareg*. We use the Jaccard similarity to create a similarity matrix for all considered pathways. As in the synthetic benchmark, we use all pathways with sizes between 50 and 500 in the biological process subtree of the Gene Ontology.

## 3 Results

First, we compare the performance of *pareg* to competing methods using a synthetic benchmark study. Second, we conduct an exploratory analysis using a breast cancer data set from TCGA.

### 3.1 Synthetic benchmark

We compare the performance of *pareg* to other enrichment tools using a synthetic data set where the ground truth is known. To do so, we select a set of activated terms and generate differential gene expression p-values using a linear model. We vary the level of noise *η* used when generating synthetic data in order to simulate different real life situations where noise can arise from measurement errors (fig. 1).

**Figure 1:**
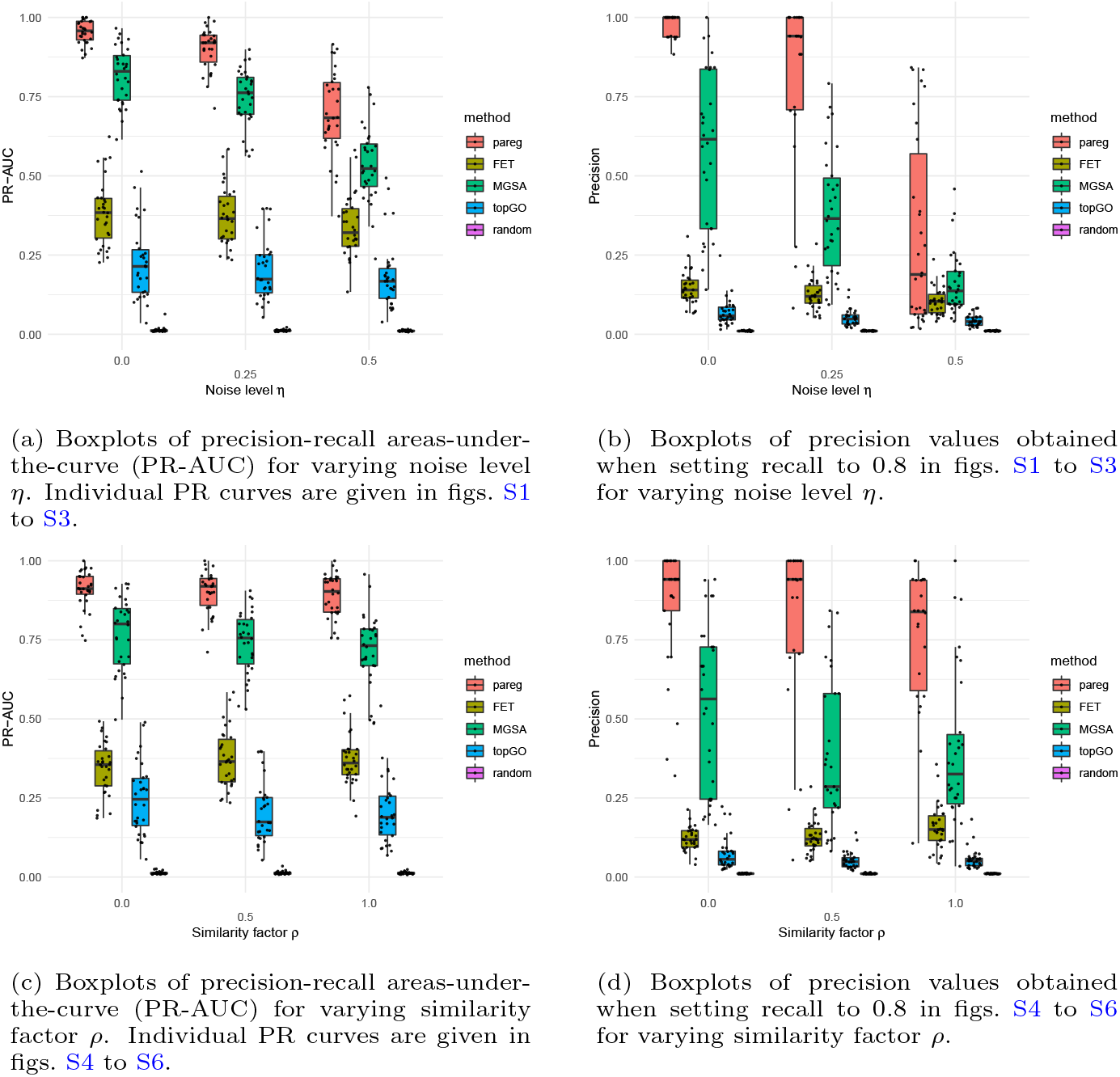
Summary of performance measures calculated for synthetic benchmark. Each point correspond to a single replicate.

In addition to *pareg*, we benchmark four other methods. MGSA is a Bayesian approach which embeds pathways in a Bayesian network and explicitly models the activation of sets of pathways (Bauer *et al*., 2010). It constitutes a modular enrichment method of competitive performance to *pareg* which does not depend on a particular pathway database. Fisher’s exact test (FET) is a classical single-term enrichment method which is still commonly used and serves as a simple alternative in the comparison (Fisher, 1922). topGO’s elim algorithm incorporates the GO tree structure into the enrichment computation and is a modular enrichment method which relies on using the Gene Ontology (Alexa *et al*., 2006). The null model serves as the baseline indicating how random guessing would perform. It assigns a random enrichment p-value between 0 and 1 to each pathway.

We observe that *pareg* consistently outperforms all competing methods over a wide range of parameter values (fig. 1). For varying levels of noise *η* = 0, 0.25, 0.5 and similarity factor *ρ* = 0, 0.5, 1, *pareg* achieves the highest mean areas under the precision-recall curve (PR-AUC) in all cases (figs. 1a and 1c). *pareg* clearly outperforms the singular enrichment method FET, which emphasizes that the proposed method of including term-term relations in the enrichment computation yields an advantage when working with large and redundant pathway databases. Out of all other benchmarked methods, MGSA performs closest to *pareg* indicating that its Bayesian model-based approach which explicitly handles term-term relations in a database-agnostic way is to some extent able to deal with the clustered pathway database. topGO performs slightly worse than FET. It explicitly uses the GO tree structure and performs successive enrichment tests which are individually similar to FET. This approach is not able to appropriately process the clustering structure assumed in the synthetic benchmark which is not based on a tree.

When increasing the noise level *η*, we observe that FET and topGO show a smaller decrease in performance than *pareg* and MGSA (fig. 1a). This is in line with the observation that the precision of FET and topGO remains nearly constant when fixing the recall (fig. 1b). For example, at a recall of 80% *pareg* has a median precision of 94% for *η* = 0.25 while MGSA has a median precision of 37%. FET and topGO have median precision values of 12% and 5% respectively. For *pareg* and MGSA, most PR-AUC is lost for large values of recall where FET and topGO show poor performance even for small *η*.

When increasing the similarity factor *ρ*, we see that *pareg* remains at roughly the same PR-AUC (fig. 1c) and only slightly decreases in precision at a fixed recall level (fig. 1d), while MGSA shows a stronger decrease in performance. For example, fixing recall to 80% at *ρ* = 0.5 yields a median precision of 94% for *pareg*. MGSA, FET and topGO have median precision values of 29%, 12% and 5% respectively. This indicates that *pareg* is better able to deal with varying levels of clustering in the set of dysregulated pathways. topGO exhibits a slight decline in performance as its tree-based approach is not able to handle the clustering structure induced by the Jaccard similarity measure. As FET does not incorporate term-term relations into the enrichment computation, we observe no dependence on *ρ*.

### 3.2 Exploratory analysis of breast cancer samples

To investigate the behavior of *pareg* on real data, we use it to run a pathway enrichment analysis on breast cancer (BRCA) samples from TCGA with terms from the Gene Ontology biological process subtree. We order the terms by their absolute enrichment level and list the top 25 results in fig. 2a as well as visualize the top 50 non-isolated results in a network (fig. 2b).

**Figure 2:**
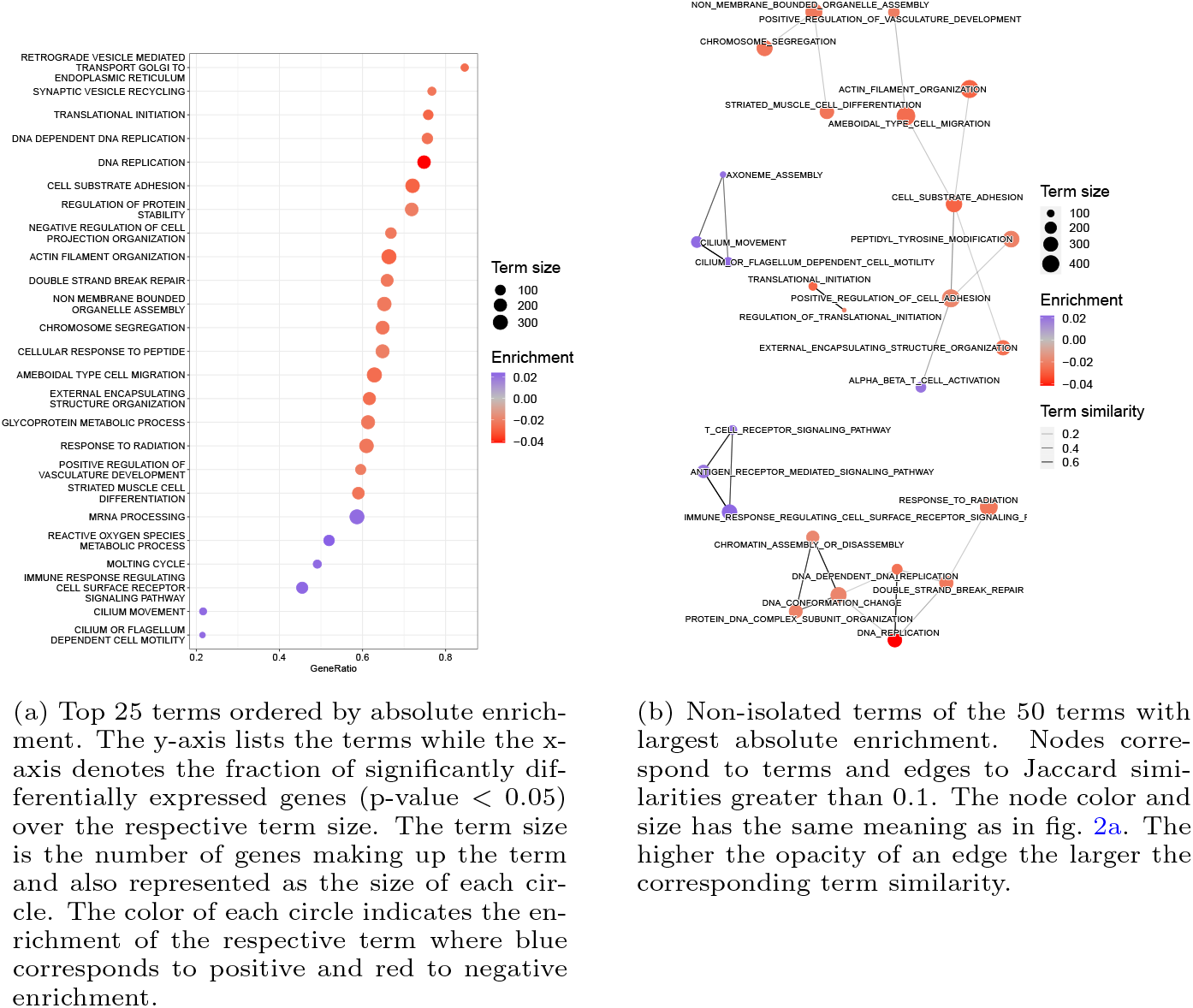
Summary of term enrichment results obtained for TCGA breast cancer samples (normal versus tumor) and the biological process subtree of the Gene Ontology.

The largest cluster of the network visualization is made up of 8 nodes and features terms related to cell migration such as ameboidal-type cell migration and actin filament organization. It has been recognized that cancer cells can use amoeboid migration as their preferred migratory strategy (Graziani *et al*., 2021). In particular, it has been shown that treatment via endocrine therapy inhibits this kind of migration in breast cancer. Furthermore, it has been shown that the organization of actin stree fibers promotes proliferation of pre-invasive breast cancer cells (Tavares *et al*., 2017). The dysregulation of cell adhesion dynamics has also been investigated in the literature (Maziveyi and Alahari, 2017) and is captured by the enrichment of the cell-substrate adhesion and positive regulation of cell adhesion terms. In addition, the peptidyl-tyrosine modification term is enriched. Tyrosine acts as a key player in the initiation of proteins to focal adhesion sites. Apart from this, the influence of tyrosine phosphatases for many different cancer types (Motiwala and Jacob, 2006) and of tyrosine kinases specifically for breast cancer (Biscardi *et al*., 2000) has been recognized.

The second largest cluster made up of 7 nodes is thematically related to DNA replication and conformation changes. These processes are of high relevance to cancers in general (Jia *et al*., 2017) as well as breast cancer specifically (Ghimire *et al*., 2020). Furthermore, the importance of double-strand break repair has been captured by the enrichment of the corresponding term (Bau *et al*., 2007).

A few smaller clusters remain. One cluster of three nodes contains the terms chromosome segregation, non-membrane-bounded organelle assembly and striated muscle cell differentiation. The importance of chromosomal stability and the impact of proteins which modulate it have been highlighted for breast cancer (Garcia and Lizcano, 2016). Furthermore, it has been observed that breast cancer cells exhibit non-random chromosome segregation (Liu *et al*., 2013). In addition, striated muscle cell differentiation has been linked to the metastatic potential of breast cancer cells (Nikulin *et al*., 2021). Another cluster of three nodes contains the terms cilium movement, cilium or flagellum-dependent cell motility and axoneme assembly. It has been shown that the expression of cilia is downregulated in various types of cancer, including breast cancer (Higgins *et al*., 2019). It furthermore has impact on the regulation of cancer development (Fabbri *et al*., 2019). The related enrichment of the axoneme assembly terms suggests the importance of the assembly and organization of an axoneme. This constitutes a novel finding and suggests further experimental investigations. The last cluster with three nodes contains the terms T-cell receptor signaling pathway, antigen receptor-mediated signaling pathway and immune response-regulating cell surface receptor signaling pathway. Both the relevance of the T-cell receptor signaling (Shah *et al*., 2021) and immune response-regulating cell surface receptor signaling term (Rezaei-Tavirani *et al*., 2019) has been recognized. The possibility of investigating the antigen receptor-mediated signaling pathway for a Chimeric antigen receptor T cell therapy has very recently been considered (Yang *et al*., 2022). Finally, the two node cluster contains the terms translational initiation and regulation of translational initiation. The regulation of translation via changed expression of the eukaryotic translation initiation factor 3 has been observed to play a positive role in breast cancer progression (Grzmil *et al*., 2010).

In addition to the network clusters, we also detect individually enriched pathways (fig. 2a). We find the retrograde vesicle-mediated transport, Golgi to endoplasmic reticulum term to be enriched. The potential implications of this apparatus have already been discussed (Spang, 2013), but have, to the best of our knowledge, not been linked to breast cancer specifically. The synaptic vesicle recycling term is also enriched. Its potential as a therapeutic target has been recognized (Li and Kavalali, 2017), however not in the context of breast cancer. In both cases, our results suggest the novel finding that these pathways may be especially relevant to breast cancer and that further experimental validations in that direction would be interesting.

We also demonstrate the effectiveness of network regularization by comparing the enrichments to results obtained from running *pareg* without the network regularization term (fig. S7) and from FET (fig. S8). In both cases, much fewer clusters are observed, making the biological interpretation more difficult. This indicates that employing the network regularization term is useful for better understanding of the enrichment results.

## 4 Discussion

We have developed a novel pathway enrichment method called *pareg* which is based on a regularized generalized linear model. It makes use of LASSO and network fusion penalty terms to produce a sparse and coherent list of enriched pathways. The network fusion term incorporates a pathway similarity network which models functional relations between pathways and clusters pathways as part of the enrichment computation in order to handle large and redundant pathway databases.

In a synthetic benchmark, we show that *pareg* is able to outperform single-term enrichment methods such as Fisher’s exact test, a popular tool explicitly including the GO tree in its calculations as well as a model-based approach which embeds pathways in a Bayesian network.

In an exploratory analysis with breast cancer samples, we are able to recover many relevant pathways already known in literature, as well as suggest novel ones which pose interesting future targets for experimental validation.

We note that *pareg* assumes that a linear combination of gene-pathway memberships is driving the overall pathway dysregulation, an assumption which may reduce the algorithm’s applicability in certain biological environments, such as, for example, the interactions between genes in myocardial infarction as measured by mRNA expression profiles (Hartmann *et al*., 2016).

Due to the flexibility of the regression approach, potential future work could go in many directions. Instead of modeling the response variable using a Beta distribution, one may use a beta-uniform mixture which has been suggested for p-values (Pounds and Morris, 2003). As the network fusion penalty depends on a general similarity matrix, different measures could be explored. For example, there exist a wide range of different semantic similarity measures which have been used to relate GO terms (Jiang and Conrath, 1997; Lin *et al*., 1998; Resnik, 1999; Schlicker *et al*., 2006; Wang *et al*., 2007; Zhao and Wang, 2018). Alternatively, similarity measures which embed sets of genes in protein-protein interaction networks and compare their localization have been shown to be useful for predicting disease status; they could be another viable choice (Bass *et al*., 2013; Menche *et al*., 2015).

Furthermore, the potential effects of other regularization terms are interesting. Using an Elastic-Net term instead of LASSO or stability selection (Meinshausen and Bühlmann, 2010) could improve the sparsity of the coefficient vector. Instead of the network fusion term, regularizations such as hierarchical feature regression (Pfitzinger, 2021), regularized k-means clustering (Sun *et al*., 2012) or group LASSO (Yuan and Lin, 2006) can be used to incorporate term-term relations and may exhibit more desirable statistical properties, such as stronger robustness to noise, smaller sample size requirements and faster convergence of the optimizer. Due to these regularization terms, it is not immediately possible to compute confidence intervals for each entry of the estimated coefficient vector. The de-biased LASSO approach (Xia *et al*., 2020) can be explored to get a better understanding of the uncertainty involved in the enrichment computation.

Finally, while there have been programming language specific efforts to standardize gene set enrichment benchmarking workflows (Geistlinger *et al*., 2021), no widely accepted consensus has been found. The benchmarking workflow we implement is written in the workflow management system Snakemake (Mölder *et al*., 2021) and thus allows easy integration of additional tools as well as reproducible execution on different back ends. We thus hope that other enrichment tools can use a similar approach to enable comparative benchmarks of new methodologies.

## 5 Data availability

The code used to construct the synthetic data sets is available as part of the R/Bioconductor software package *pareg*. The experimental data used in the exploratory analysis is available as GSE62944 on the Gene Expression Omnibus. The pathway database has been obtained from the Gene Ontology resource.

## 6 Code availability

The method *pareg* is freely available as an R package on Bioconductor (https://bioconductor.org/packages/release/bioc/html/pareg.html) as well as on https://github.com/cbg-ethz/pareg. The GitHub repository also contains the Snakemake (Mölder *et al*., 2021) workflows needed to reproduce all results presented here.

