## Supplementary figures for "Coherent pathway enrichment estimation by modeling inter-pathway dependencies using regularized regression"

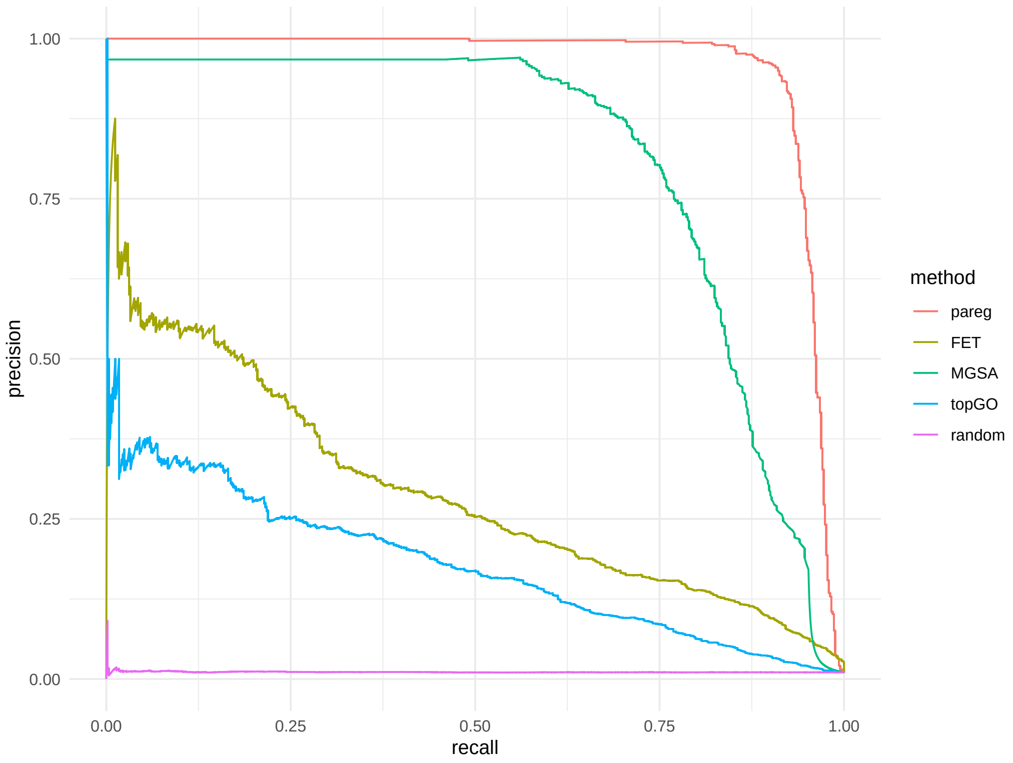


Figure S1: Precision-Recall (PR) curves aggregated over all replicates for noise level $\eta = 0$.

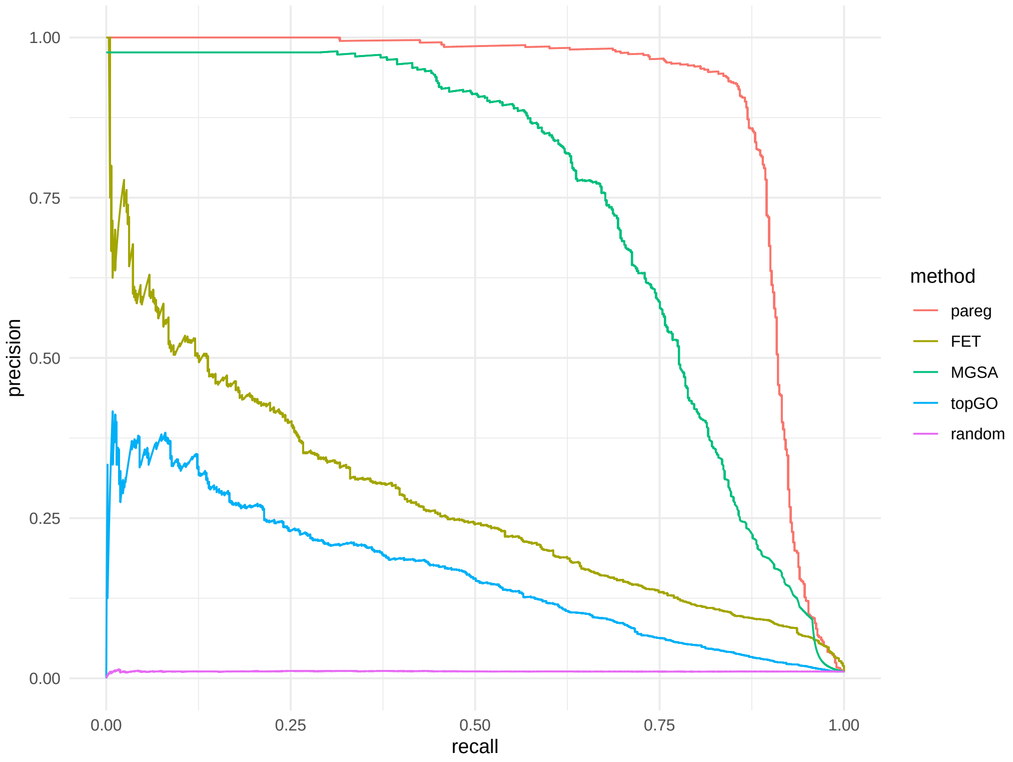


Figure S2: Precision-Recall (PR) curves aggregated over all replicates for noise level $\eta = 0.25$.

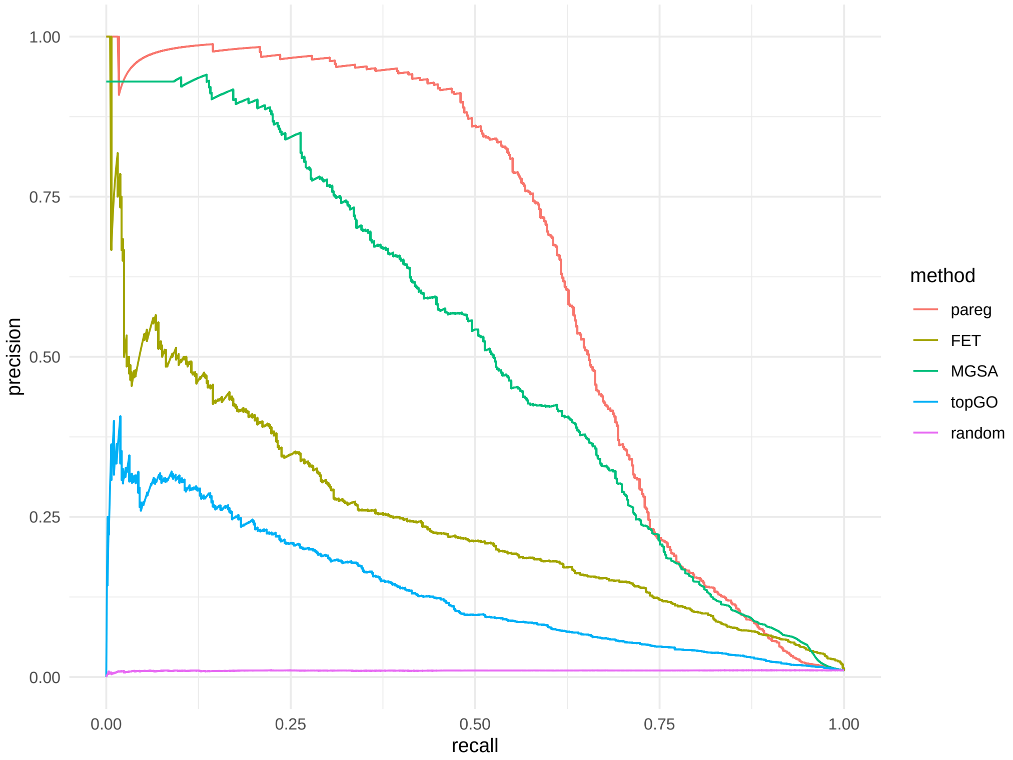


Figure S3: Precision-Recall (PR) curves aggregated over all replicates for noise level $\eta = 0.5$.

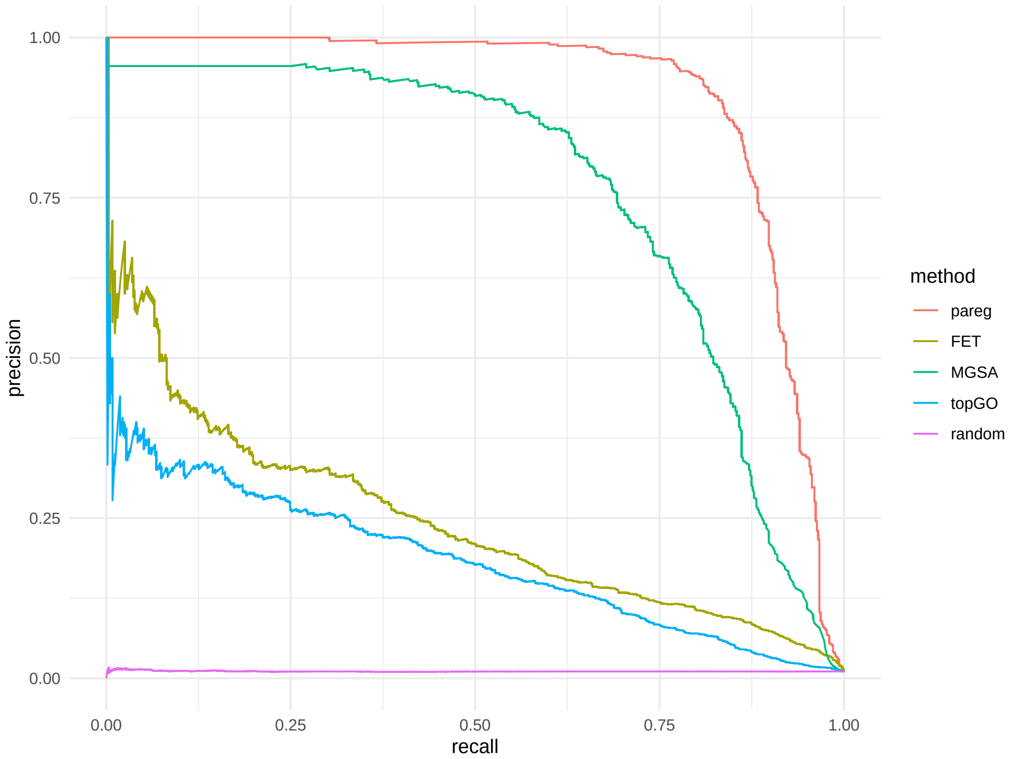


Figure S4: Precision-Recall (PR) curves aggregated over all replicates for similarity factor $\rho = 0$.

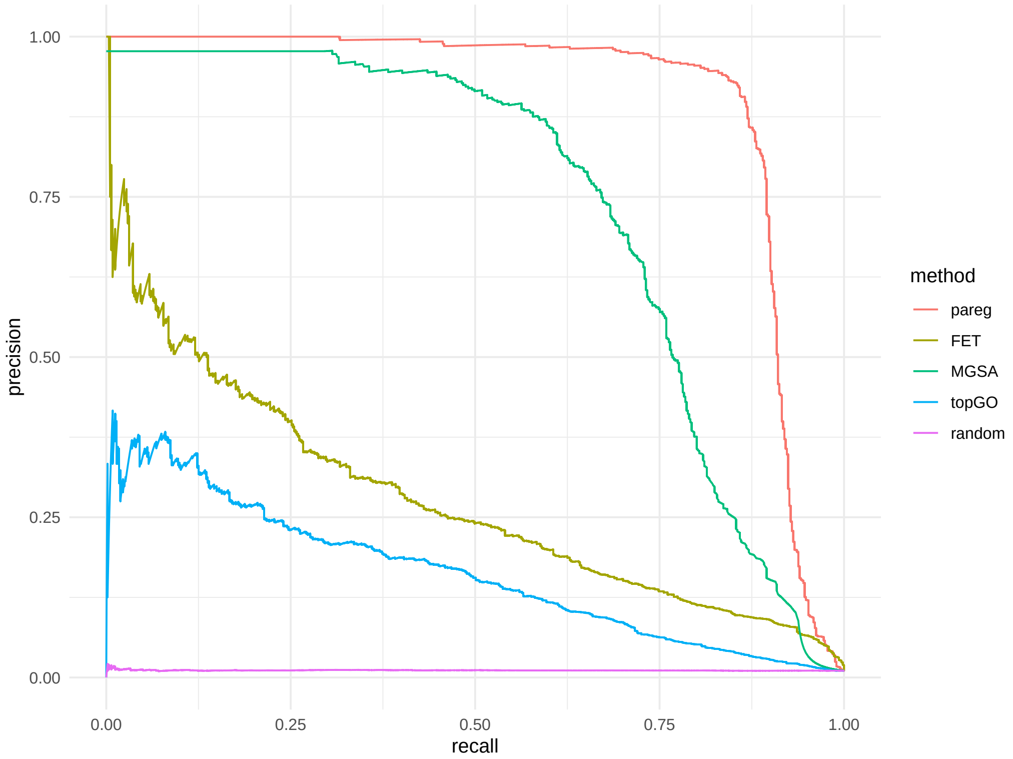


Figure S5: Precision-Recall (PR) curves aggregated over all replicates for similarity factor $\rho = 0.5$.

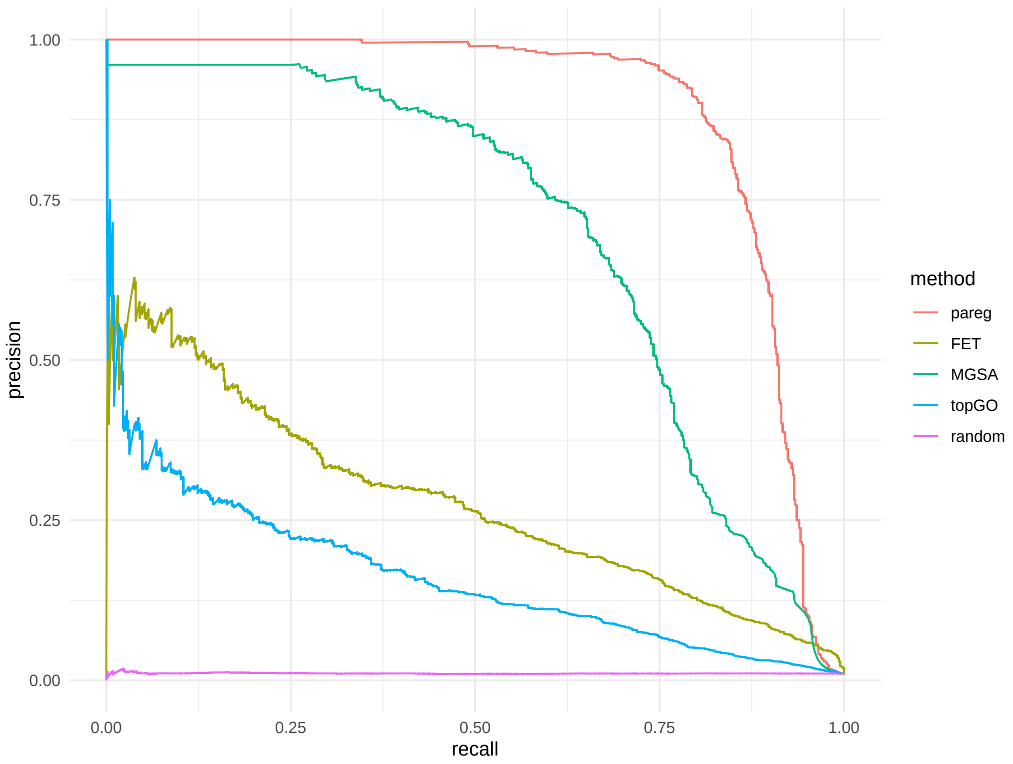


Figure S6: Precision-Recall (PR) curves aggregated over all replicates for similarity factor $\rho = 1$.

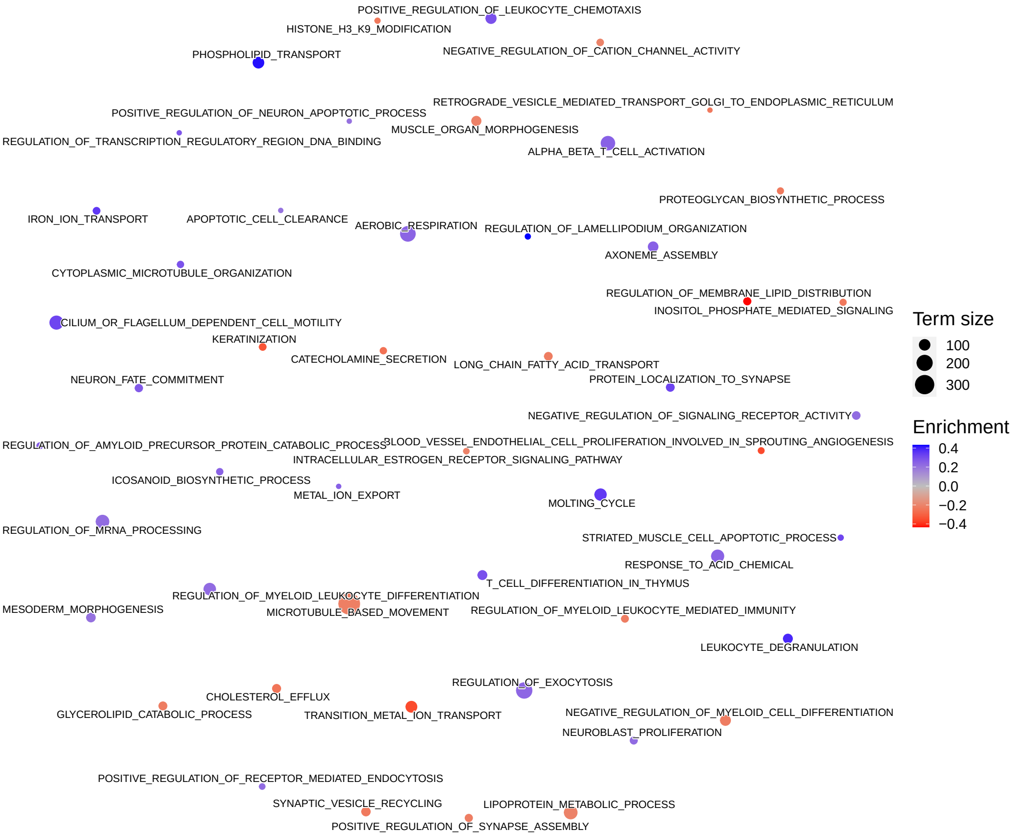


Figure S7: Term network for *pareg* without network regularization with same parameters as in fig. 2b) except for also including isolated terms.

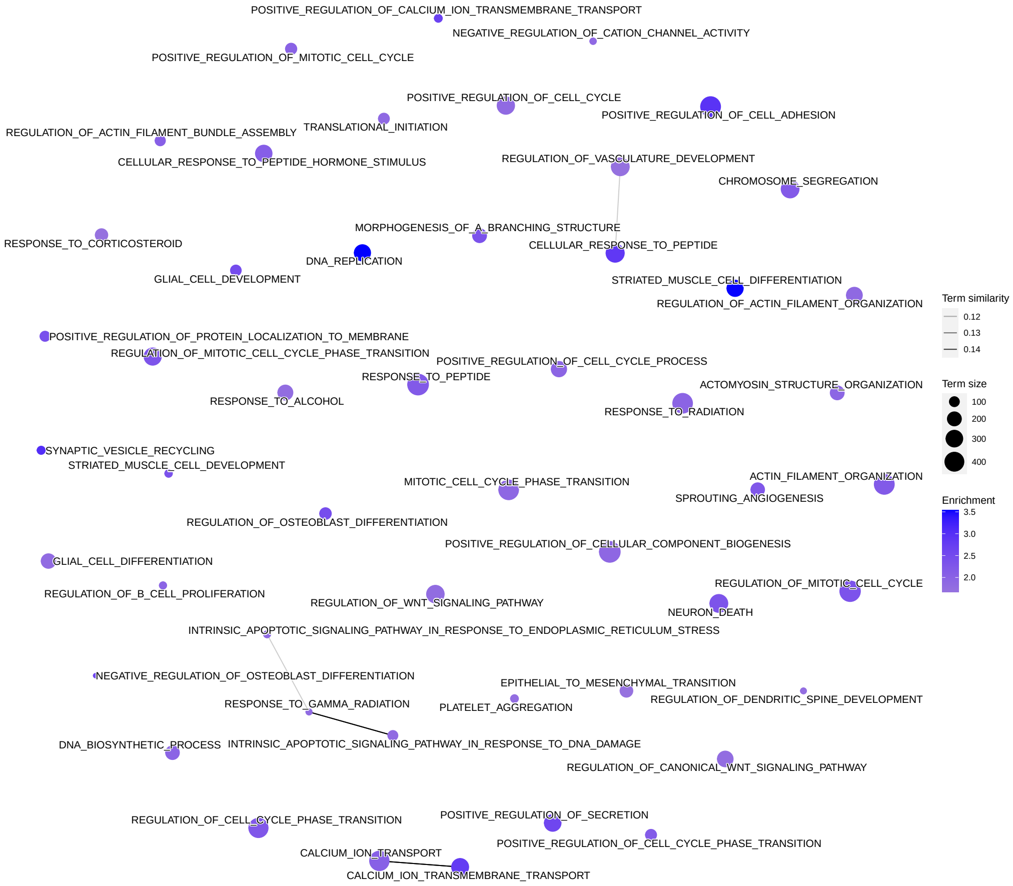


Figure S8: Term network for FET with same parameters as in fig. 2b) except for also including isolated terms. The enrichment score is the negative decadic logarithm of the p-value.
